# Self-assembly of DNA nanostructures in different cations

**DOI:** 10.1101/2023.05.04.539416

**Authors:** Arlin Rodriguez, Dhanush Gandavadi, Johnsi Mathivanan, Tingjie Song, Bharath Raj Madhanagopal, Hannah Talbot, Jia Sheng, Xing Wang, Arun Richard Chandrasekaran

## Abstract

The programmable nature of DNA allows the construction of custom-designed static and dynamic nanostructures, and assembly conditions typically require high concentrations of magnesium ions which restricts their applications. In other solution conditions tested for DNA nanostructure assembly, only a limited set of divalent and monovalent ions have been used so far (typically Mg^2+^ and Na^+^). Here, we investigate the assembly of DNA nanostructures in a wide variety of ions using nanostructures of different sizes: a double-crossover motif (76 bp), a three-point-star motif (∼134 bp), a DNA tetrahedron (534 bp) and a DNA origami triangle (7221 bp). We show successful assembly of a majority of these structures in Ca^2+^, Ba^2+^, Na^+^, K^+^ and Li^+^ and provide quantified assembly yields using gel electrophoresis and visual confirmation of a DNA origami triangle using atomic force microscopy. We further show that structures assembled in monovalent ions (Na^+^, K^+^ and Li^+^) exhibit up to a 10-fold higher nuclease resistance compared to those assembled in divalent ions (Mg^2+^, Ca^2+^ and Ba^2+^). Our work presents new assembly conditions for a wide range of DNA nanostructures with enhanced biostability.

## 1. INTRODUCTION

DNA has become an attractive material for the assembly of nanostructures with custom-designed shapes, high size homogeneity, addressable features and capability for stimuli-responsive reconfiguration.^1–3^ Aided by recent advancements in DNA synthesis,^4^ programmed assembly^5^ and chemical functionalization strategies,^6^ DNA nanostructures are used in applications including diagnostics,^7^ drug delivery,^8^ rewritable data storage,^9^ molecular electronics,^10^ neural networking,^11^ and single molecule biophysics.^12^ Assembly of DNA nanostructures is typically achieved through the cooperative assembly of short DNA strands,^13^ modular assembly of DNA motifs,^14^ hierarchical assembly of DNA tiles into larger structures^15^ or the DNA origami strategy.^16^ While the capacity of DNA-based self-assembly has been expanded into the micrometer scale^17^ and gigadalton size,^18^ most methods still require magnesium-containing buffers for DNA self-assembly. Magnesium is widely considered as an essential component of DNA self-assembly for its role in screening the inter-helical repulsion^19^ and stabilizing the stacked form of branched DNA junctions.^20^ Despite this critical role in DNA self-assembly, magnesium ions can sometimes have adverse effects by causing aggregation of DNA-nanoparticle complexes at high ionic concentrations,^21^ enhancing nuclease activity,^22^ affecting mineralization of DNA origami structures,^23^ interfering with drug loading due to metal complexation of small molecule drugs^24^ or by modulating intercalative properties,^25^ and by affecting reconfiguration of pH responsive DNA nanostructures.^26^ Further, some applications may require a different ion for DNA nanostructure assembly such as to stabilize proteins arranged on DNA nanostructures,^27^ for cation-responsive reconfiguration^28^ and metal-mediated base pairing.^29^

To mitigate the requirement of magnesium, recent studies have shown the assembly of representative DNA origami structures (triangular and multi-helix bundles) in low-magnesium buffers^30^ and in buffers containing Na^+^.^31^ Cation-free assembly of DNA nanostructures has also been accomplished using ethylenediamine buffer, where the protonated forms of ethylenediamine replaces the need for divalent cations such as Mg^2+^.^22^ Beyond these studies, the effect of other metal ions on the assembly of DNA nanostructures have not been explored in detail. Expanding the choice of ions for DNA nanostructure assembly would be useful in improving co-assembly with nanoparticles,^32^ to control the attachment of DNA nanostructures to lipid membranes,^33^ to modulate the activity of Mg^2+^-dependent enzymes^34^ and in guiding DNA nanostructure assembly by cation-mediated DNA-DNA attraction.^35^

In this work, we investigated the effect of different monovalent and divalent cations on the assembly of DNA nanostructures ranging in size from tens to thousands of base pairs. As model DNA nanostructures, we chose the double crossover (DX) DNA motif (76 bp) constructed using cooperative assembly of four component strands, a symmetric three-point-star motif (∼134 bp) assembled using three unique strands, a DNA tetrahedron (534 bp) hierarchically assembled from three-point-star motifs and a triangular DNA origami structure (7221 bp). We characterized DNA nanostructure assembly and quantified the assembly yields in twelve different cations using gel electrophoresis and provide visual confirmation of DNA origami assembly using atomic force microscopy (AFM). We also determined the biostability of these structures against DNase I and in fetal bovine serum (FBS) and show that the choice of cations for DNA nanostructure assembly can play a significant role in their enhanced biostability.

## 2. RESULTS

### 2.1 Assembly of double crossover (DX) DNA motif

To demonstrate the assembly of DNA nanostructures using different metal ions, we first chose the DX motif, a structure containing two adjacent double helical domains connected by two crossover points (**Figure 1a-b** and **Figure S1**).^36^ DX motifs have been used in the assembly of 2D lattices and is a part of larger structures such as DNA origami that involve multiple DNA crossovers. We used a DX motif composed of four DNA strands with 16 base pairs between the crossover points. We assembled the DX motif in tris-acetate-EDTA (TAE) buffer containing 12.5 mM Mg^2+^ (typical annealing buffer used for DNA motifs) and validated proper assembly using non-denaturing polyacrylamide gel electrophoresis (PAGE) (**Figure 1c**).

**Figure 1.**
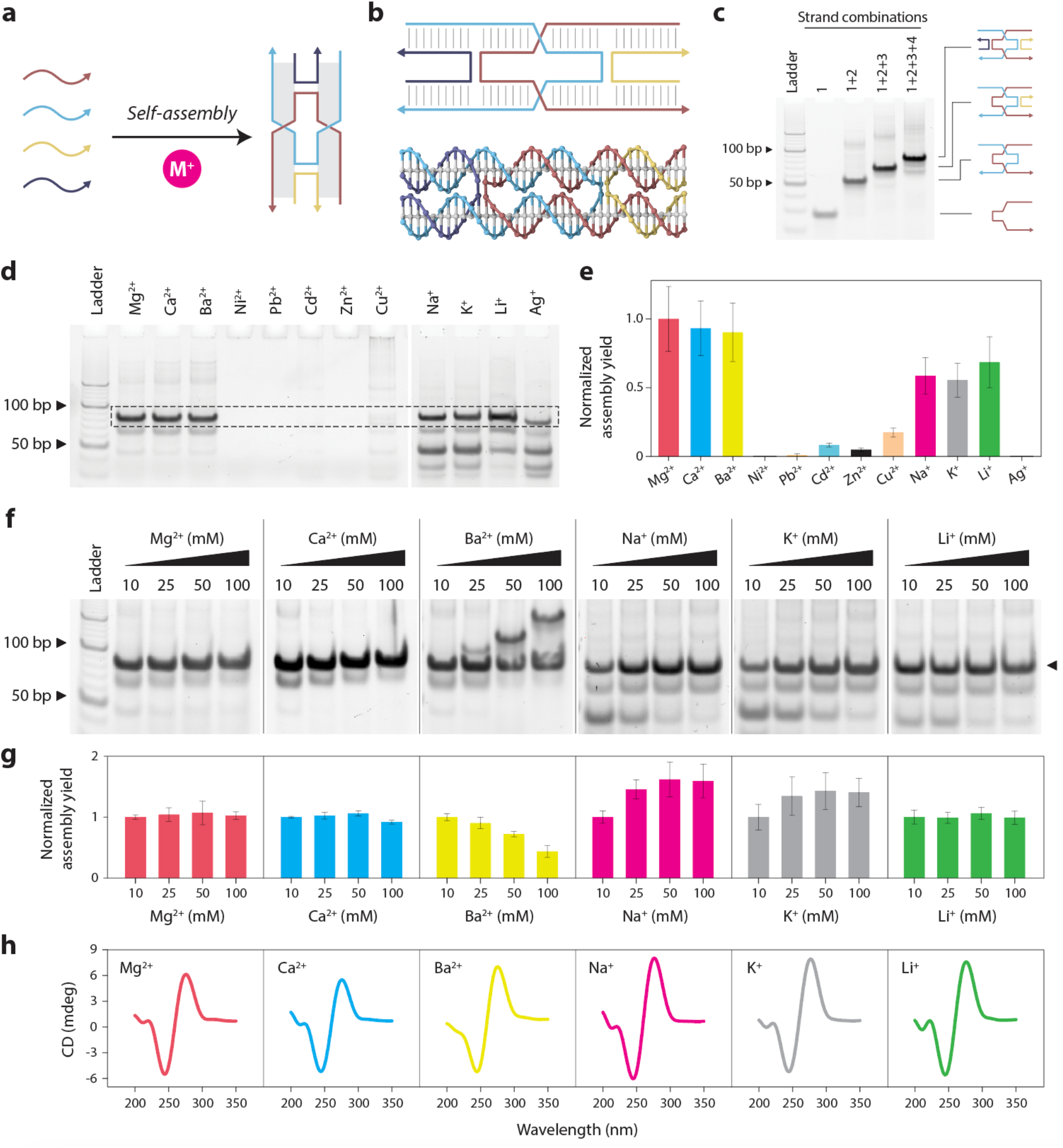
Assembly and characterization of the DX motif in different cations. (a) Schematic showing assembly of DX motif using multiple component strands in buffer containing metal ions (M^2+^). (b) Illustration and molecular model of a DX motif. (c) Non-denaturing PAGE showing assembly of the DX motif in a typical Mg^2+^-containing buffer. (d) Non-denaturing PAGE showing assembly of DX motif in different divalent and monovalent ions. (e) Quantified results from (d) showing the assembly yield of DX motif. (f) Non-denaturing PAGE analysis of DX motif assembly in buffer containing different concentrations (10, 25, 50, 100 mM) of Mg^2+^,Ca^2+^,Ba^2+^,Na^+^, K^+^and Li^+^.(g) Quantified results from (f) showing the assembly yield of DX motif in different ion concentrations. Assembly yields are normalized to the yield in 10 mM ion for each case. (h) CD spectra of DX motif assembled in 10 mM ions. Data represent mean and error propagated from standard deviations of experiments performed in triplicates.

To test DX assembly in different cations, we replaced the Mg^2+^ in the TAE buffer with different divalent (Ca^2+^, Ba^2+^, Ni^2+^, Cd^2+^, Pb^2+^, Zn^2+^ and Cu^2+^) and monovalent (Na^+^, K^+^, Li^+^ and Ag^+^) ions. We annealed the structure and characterized the assembly of DNA complexes using non-denaturing PAGE (**Figure 1d**). For these experiments, we performed PAGE analysis using running buffer that did not contain Mg^2+^ so as to reduce any effect it may have on DNA nanostructure analysis. Results showed that different ions had different effects on DNA self-assembly, with some ions interfering with the assembly process entirely, resulting in no formation of the desired complex. The DX motif was assembled properly in Ca^2+^, Ba^2+^, Na^+^, K^+^, and Li^+^, showing a band on the gel similar to that of the DX assembled in Mg^2+^. We obtained the assembly yield in each case by quantifying the band on the gel corresponding to the DX structure (**Figure 1e**).

We then annealed the DX structure in different concentrations of each cation to test whether assembly is affected by different metal ion concentrations (**Figure 1f** and **Figure S2**). We performed the experiments in triplicates and obtained the assembly yield in buffers containing 10, 25, 50, and 100 mM ions (**Figure 1g**). For the DX motifs assembled in Mg^2+^, Ca^2+^, and Li^+^, assembly yields remained similar with increasing ion concentration whereas for structures assembled in Na^2+^ and K^+^, assembly yield increased with ion concentration. For Ba^2+^, we observed the appearance of bands corresponding to higher order structures at ion concentrations above 25 mM, and the yield of the DX motif reduced with increasing ion concentration. Our observation of higher order assemblies with increasing Ba^2+^ concentrations could be related to the role of Ba^2+^ in binding DNA junctions^37^ and the formation of G-quadruplexes in specific DNA sequences.^38^ Assembly of DX was not observed in samples with Ni^2+^, Cd^2+^, Pb^2+^, Zn^2+^, Cu^2+^ and Ag^+^, indicating degradation or aggregation of the DNA strands or structures. Our results are consistent with the known interactions of these different cations with nucleic acids. Alkali metals (eg: Li^+^, Na^+^ and K^+^) and alkaline earth metals (eg: Mg^2+^, Ca^2+^ and Ba^2+^) mainly interact with the phosphate groups on the backbone and can thus stabilize the DNA structure.^39^ On the other hand, transition metals (eg: Ni^2+^, Cd^2+^, Pb^2+^, Zn^2+^, and Cu^2+^ used in this study) interact with the nucleobases and may destabilize the duplex structure of DNA,^40^ and thus also affect the assembly of DNA nanostructures.

For the conditions we observed assembly of the DX motif, we then performed circular dichroism (CD) and UV melting studies. CD spectra of the DX motifs assembled in buffer containing 10 mM Ca^2+^, Ba^2+^, Na^+^, K^+^ and Li^+^ were similar to that of the DX motif assembled with Mg^2+^, indicating that the assembly was not affected when Mg^2+^ was replaced by these ions (**Figure 1h**). The CD spectra were also consistent with the spectrum we reported earlier for a DX motif,^41^ indicating that the underlying structure was B-form DNA in all these conditions. UV melting studies showed that the melting temperature (Tm) for DX motif assembled in TAE-Mg^2+^ was 69 °C while the Tm for the motifs assembled in 10 mM Ca^2+^, Ba^2+^, Na^+^, K^+^, and Li^+^ were 65, 63, 48, 47, and 49 °C, respectively (**Figure S3**)

### 2.2 Assembly of the three-point star motif and DNA tetrahedron

We next investigated whether the different cations that worked for DX self-assembly could also be used for other DNA nanostructures. To demonstrate this, we chose a DNA tetrahedron hierarchically self-assembled from three-point-star motifs (**Figure 2a**).^15^ This model system serves two purposes: (1) to study the effect of different cations in another DNA motif (the three-point-star) and (2) to study the effect of different cations in sticky end cohesion (formation of DNA tetrahedron from three-point-star motifs). The three-point-star motif is assembled from three unique strands: a 78 nt long (L) strand, a 42 nt medium (M) strand and a 21 nt short (S) strand. The motif contains three arms, each of which consists of two double helical domains connected by a single crossover. The motif contains 5T loops in the middle to provide flexibility to assemble into a DNA tetrahedron. Four units of the three-point-star motif assemble via sticky end cohesion to form a DNA tetrahedron with six edges and four faces (**Figure 2a**). To distinguish assembly of the individual motif and the tetrahedron, we designed a three-point-star motif without sticky ends to prevent assembly into the DNA tetrahedron (**Figure S4**) and validated assembly of the structures using non-denaturing PAGE (**Figure 2b**).

**Figure 2.**
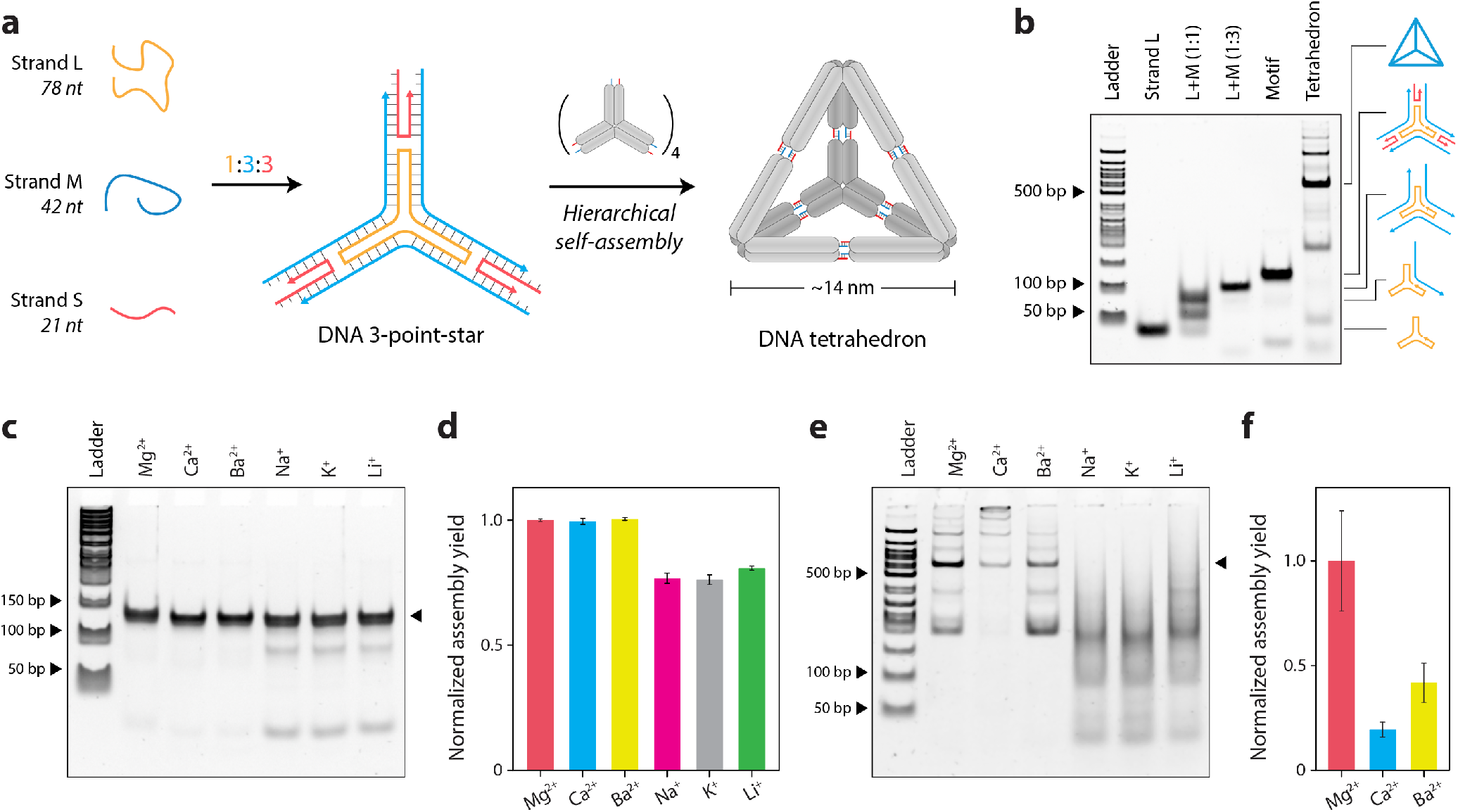
Assembly and characterization of three-point-star and DNA tetrahedron in different cations. (**a**) Schematic showing the assembly of the three-point-star motif and further hierarchical assembly into a DNA tetrahedron. (**b**) Non-denaturing PAGE showing assembly of the three-point-star motif and DNA tetrahedron. (**c**) Non-denaturing PAGE and (**d**) quantified assembly yields of three-point-star in different ions. (**e**) Non-denaturing PAGE and (**f**) quantified assembly yields of the DNA tetrahedron in different ions. Data represent mean and error propagated from standard deviations of experiments performed in triplicates.

We annealed the blunt-ended three-point-star motif in TAE buffer containing different cations (at 10 mM concentration) and tested the assembled structures using non-denaturing PAGE (**Figure 2c**). Gels run in buffer without Mg^2+^ showed a split band while gels run in buffer containing Mg^2+^ showing proper bands corresponding to the structure (**Figure S5**). Since the three-point-star motif consists of two double helical domains per arm as well as 15 unpaired bases at the center (three 5T loops), it is possible for the structure to exist in different angular conformations, thus showing split bands on a gel in the absence of magnesium. Mg^2+^ is known to reduce the conformational entropy and rigidify DNA structures,^42^ causing the motif to be more compact and run as a single discrete band in running buffer containing Mg^2+^. We ran triplicates of the three-point-star assembled in different ions in buffer containing Mg^2+^ and quantified the assembly yields. We observed similar assembly yields for the three-point-star motif in 10 mM Mg^2+^, Ca^2+^, and Ba^2+^ and comparatively lower assembly yields and formation of intermediate structures in Na^+^, K^+^, and Li^+^ (**Figure 2c-d**).

Next, we used a three-point-star motif containing sticky ends to assemble the DNA tetrahedron with different cations and analyzed assembly using non-denaturing PAGE (**Figure 2e**). Although 10 mM monovalent ions could be used to form the three-point-star motifs, their subsequent assembly into the tetrahedron was highly impaired, with only Mg^2+^, Ca^2+^, and Ba^2+^ showing assembly of tetrahedra at the ion concentrations tested (**Figure 2f**). The tetrahedron assembly yields were also considerably lower in Ca^2+^ and Ba^2+^ compared to the structure assembled in Mg^2+^, indicating that while some divalent ions can yield proper assembly of individual three-point-star motifs, their efficiency in stabilizing sticky end cohesion (for tetrahedra formation) can be vastly different. We attribute these differences to the many factors involved in the hierarchical assembly of DNA motifs into larger structures, such as sticky end length and sequences (eg: GC content) as well as the concentration and type of counter ions. Only specific solution conditions allow reversible error-correcting assembly of multiple DNA motifs such as the three-point-stars into the DNA tetrahedron. While we did not observe tetrahedra formation at 10 mM Na^+^, K^+^, and Li^+^, another recent work showed that similar DNA tetrahedra can be assembled in buffer containing 100-600 mM Na^+^ or 400 mM K^+^ ions.^43^

### 2.3 Assembly of a DNA origami triangle

To further investigate the impact of different cations on the self-assembly of DNA nanostructures, we tested a larger nanostructure assembled using the DNA origami strategy.^16^ We assembled a triangle-shaped DNA origami nanostructure by folding an M13mp18 scaffold DNA (7249 nt) with 208 staple strands in TAE buffer containing 10 mM cations using a thermal annealing step (**Figure 3a**). We first characterized the formation of the DNA origami triangle using non-denaturing agarose gel electrophoresis (**Figure 3b-c**). We observed that some ions yielded proper assembly (Mg^2+^, Ca^2+^, and Ba^2+^) while other ions did not result in the formation of the desired triangle structure. DNA origami structures annealed in buffers containing Cd^2+^, Zn^2+^, Na^+^, K^+^, and Li^+^ ions showed products with slower mobility indicating unfolded structures or aggregates (**Figure 3b**). Structures assembled in Cu^2+^ and Ni^+^ showed a smear, indicating potential DNA cleavage reported earlier in these ions.^44,45^

**Figure 3.**
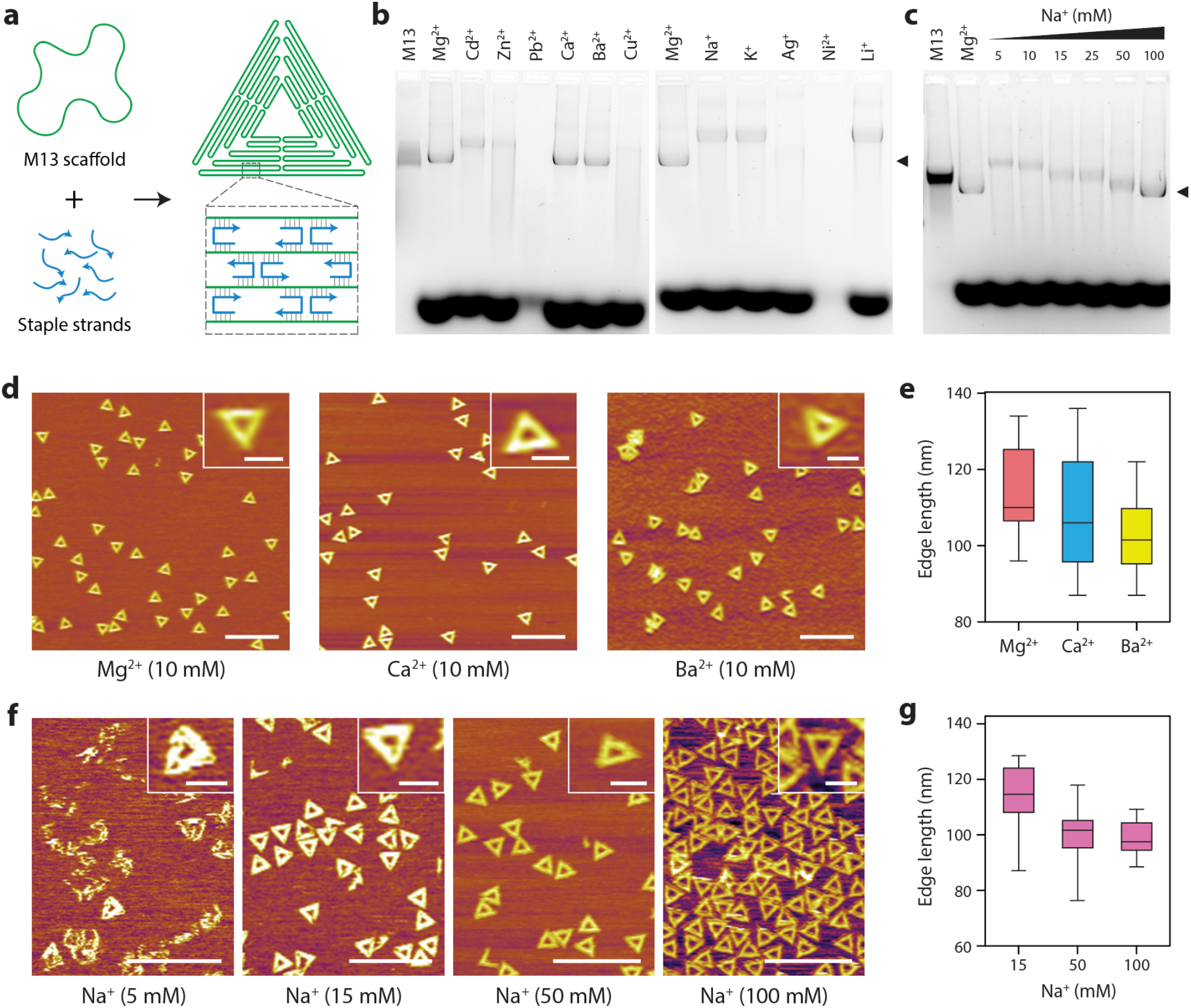
Assembly and characterization of DNA origami triangle in different cations. (a) Schematic of DNA origami assembly. (b) Agarose gel electrophoresis showing assembly of the DNA origami triangle in buffers containing different divalent and monovalent ions. (c) Assembly of the DNA origami triangle in different concentrations of Na^+^,. (d) AFM images of DNA origami triangle assembled in Mg^2+^, Ca^2+^,and Ba^2+^,ions. (e) Edge lengths of DNA origami triangles assembled in Mg^2+^, Ca^2+^,and Ba^2+^,ions. (f) AFM images of DNA origami triangle assembled in 5, 15, 50 and 100 mM Na^+^, ions. (g) Edge lengths of DNA origami triangles assembled in 15, 50 and 100 mM Na^2+^, ions. Scale bars in zoomed out images in (d) and (f) are 500 nm. Scale bars in insets are 100 nm.

We then used AFM to visually examine the formation of the DNA origami triangle in the conditions that showed proper assembly on agarose gels. AFM analysis confirmed proper assembly of the DNA origami triangle in 10 mM Ca^2+^ and Ba^2+^ ions compared to the control structures assembled in 10 mM Mg^2+^ (**Figure 3d**). We measured the triangle edge lengths in cases where we observed proper assembly and found values of 119.8 ± 4.7 nm for structures assembled in Mg^2+^ (n = 15), 109.4 ± 5.1 nm for Ca^2+^ (n = 15) and 103.4 ± 6.4 nm for Ba^2+^ (n = 15) (**Figure 3e**). It is interesting that while the size of the assemblies follows the order Mg^2+^>Ca^2+^> Ba^2+^, the actual size of the ions is Ba^2+^ > Ca^2+^ > Mg^2+^. It might appear that larger metal ions result in more compact assemblies, a trend observed in prior AFM studies of DNA in different ions.^46,47^ AFM analysis of structures assembled in other divalent ions (at 10 mM) did not yield any visible structures, while for monovalent ions, we observed proper formation only in Na^+^ (**Figure S6**). Based on a previous study that reported DNA origami assembly in high concentrations of Na^+^, we investigated DNA origami triangle assembly with increasing concentrations (5-100 mM) of Na^+^ and observed higher assembly yields with higher Na^+^ ion concentrations (**Figure 3f**). These results are also validated by our gel studies that showed compact structures corresponding to the DNA origami triangle in 50 and 100 mM Na^+^ (**Figure 3c**). We measured the triangle edge lengths for structures assembled in Na^+^ and obtained values of 113.2 ± 11.6 (n = 20) for 15 mM, 99.9 ± 8.8 nm (n = 20) for 50 mM and 98.6 ± 5.8 nm (n = 20) for the 100 mM Na^+^ condition (**Figure 3g**). The increased assembly yields we observe in higher concentrations of Na^+^ is consistent with a previous study31 that used DNA origami multi-helix bundles. The DNA origami structures reported in that study required 200 mM to 1.6 M of Na^+^ for assembly, possibly due to the packing of adjacent double helical domains into multilayer objects compared to our single-layer, two-dimensional origami structure which requires <100 mM Na^+^ for proper assembly.

### 2.4 Analysis of DNA nanostructure biostability

Next, we analyzed the biostability of DNA nanostructures assembled in different cations using a gel-based method we reported earlier (**Figure 4a**).^48,49^ We treated the DX motif with the common endonuclease DNase I for different time periods, ran the DNase I treated samples on a non-denaturing gel and quantified the band corresponding to the structure at each time point to obtain nuclease degradation profiles (**Figure 4b-c** and **Figure S7**). We observed that structures assembled in Mg^2+^ and Ca^2+^ degraded quickly (>95% degraded in 16 min), which was not surprising since DNase I is known to require Mg^2+^ or Ca^2+^ as cofactors for its enzyme activity.^50^ DX motif assembled in Ba^2+^ showed a similar degradation profile, possibly due to the preference of the nuclease for divalent cations for its activity.^51^ However, structures assembled in monovalent ions (Na^+^, K^+^, and Li^+^) were ∼40-50% intact even after 1 hour of DNase I treatment, despite the samples containing DNase I buffer that includes Mg^2+^ and Ca^2+^. Using the time constants of the degradation profiles (**Figure S8** and **Table S1**), we calculated the biostability enhancement factor (BioEF), a metric we established previously for other DNA motifs,^49^ for the DX motif assembled in these six metal ions. Compared to the structure assembled in Mg^2+^, the structures assembled in monovalent ions Na^+^, K^+^ and Li^+^ showed up to 10-fold enhanced biostability (**Figure 4d**). The reduced degradation of the DX motif in Na^+^, K^+^, and Li^+^ could be due to the inhibition of DNase I activity by monovalent ions^52^ and the different levels of thermodynamic stability offered by these cations for DNA nanostructures.^53^

**Figure 4.**
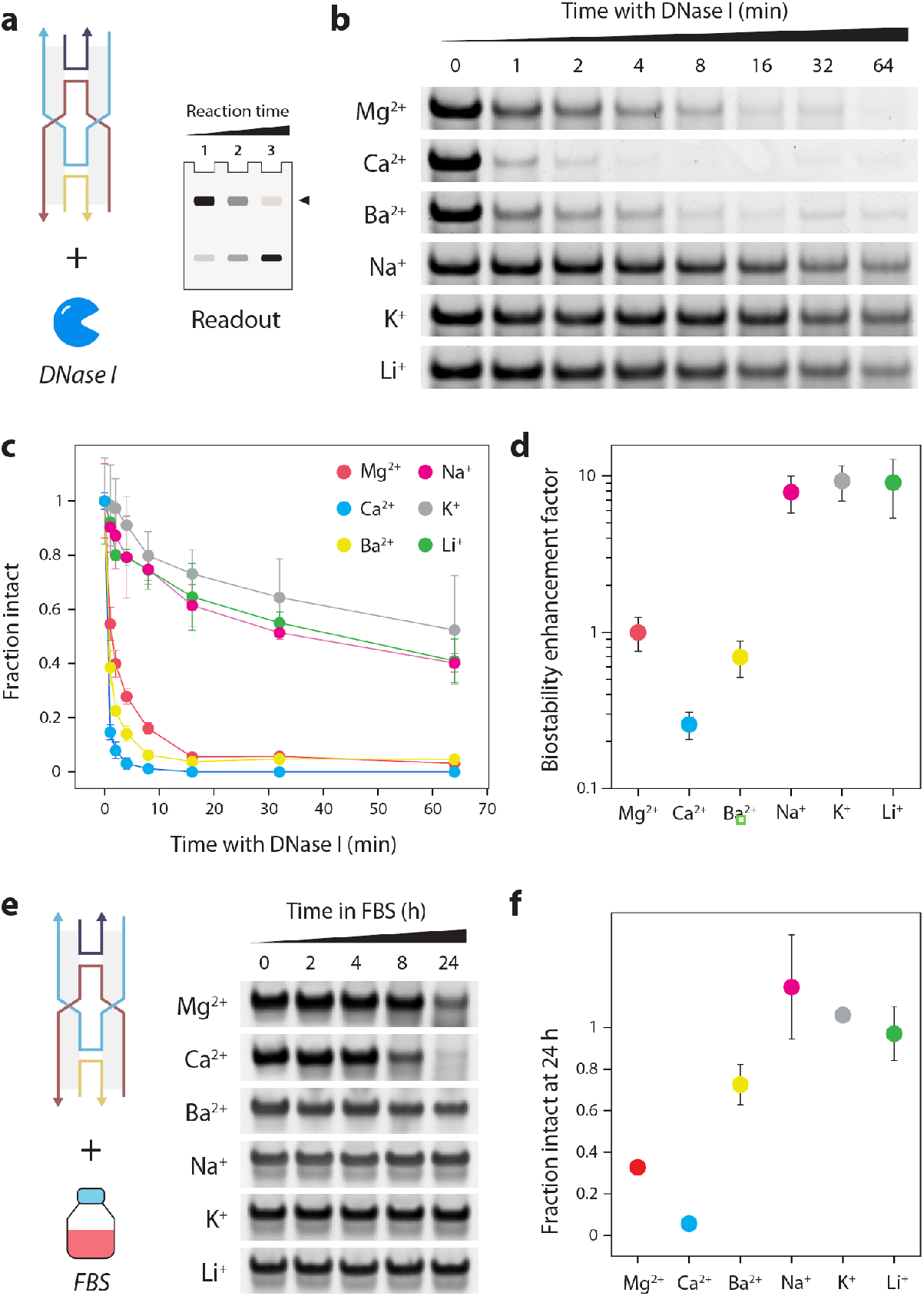
Nuclease resistance analysis of DX motif assembled in different cations. (a) Gel-based analysis of nuclease degradation. (b) Gel images showing degradation of DX motif when treated with DNase I. (c) Fraction of intact structure at different time points after treatment with DNase I. (d) Biostability enhancement factor (BioEF) calculated from time constants of nuclease degradation profiles. (e) Gel images showing degradation of DX motif in 10% FBS. (f) Fraction of intact structure after 24-hour incubation with 10% FBS. Data represent mean and error propagated from standard deviations of experiments performed with a minimum of two replicates.

To establish biological relevance, we then analyzed the biostability of the structures in FBS. We first chose the DX motif and confirmed that the structure is stable at the physiological temperature of 37 °C for 24 hours (**Figure S9**). We then incubated the DX motif in 10% FBS for different time points up to 24 hours and analyzed the samples using non-denaturing PAGE (**Figure 4e** and **Figure S10**). We quantified the intact fraction of the DX motif and observed that DX motif assembled in monovalent ions (Na^+^, K^+^ and Li^+^) showed minimal degradation even after 24 hours compared to structures assembled in divalent ions Mg^2+^ and Ca^2+^ (70-100% degraded), a trend consistent with our results in DNase I (**Figure 4f**). One difference we observed was that the DX assembled in Ba^2+^ showed similar levels of degradation to those in Mg^2+^ and Ca^2+^ when tested against DNase I, but showed a higher stability in FBS (∼30% degraded after 24 hours), possibly due to the different levels of the nuclease activity in bodily fluids. However, the overall biostability trends in both DNase I and FBS were similar, with structures assembled in monovalent ions being more biostable than those assembled in divalent ions. For the larger DNA origami triangle, structures assembled in different cations were all intact even after 24 hours of incubation in 10% FBS (**Figure S11**). Such large structures with closed packed helices are known to be more stable in serum compared to wireframe structures.^54^ Overall, our results show that assembly conditions can be a major factor in determining the biostability properties of DNA nanostructures, and assembly in monovalent ions could confer higher resistance to degradation by nucleases such as DNase I and enhanced stability in biofluids.

## 3. DISCUSSION

In this work, we have presented new results on DNA nanostructure self-assembly in the presence of different monovalent and divalent cations. Our work complements existing studies on magnesium-free assembly of DNA nanostructures (that only use Na^+^)^31,43^ and provides assembly conditions in a wider variety of cations. For the smaller DX motif, the structure has previously been assembled in 125 mM Ca^2+^ to template morphology changes in calcium carbonate,^55^ while here we show that the structure can be assembled in 10 mM Ca^2+^. For the DNA tetrahedron assembled through sticky end cohesion, we analyzed assembly at 10 mM ion concentrations and did not observe assembly in monovalent ions. A recent study that was published during our work showed that similar DNA tetrahedra can be assembled at higher Na^+^ (100-600 mM) and K^+^ (400 mM) concentrations.^43^ To our knowledge, our study is the first to show assembly of a variety of DNA nanostructures in Ba^2+^, with assembly yields comparable to a Mg^2+^-containing buffer for the DX and three-point-star motifs as well as for the larger DNA origami triangle. While our work shows that a variety of DNA nanostructures can be assembled in different cations, the choice of cation would be dependent on the specific design of the structure and the application.^30^ For example, nanostructure design involving non-canonical structures (eg: i-motifs) may be destabilized in the presence of monovalent ions (Na^+^, K^+^ and Li^+^).^56^

Prior work by other groups has demonstrated several strategies for substituting Mg^2+^ in buffers post-assembly as well as using other ions in combinations with Mg^2+^. For example, surface-assisted growth of DNA origami arrays from individual origami units assembled in Mg^2+^ has been achieved using monovalent ions such as Na^+^, K^+^, and Li^+ 57^ or by adjusting the relative concentrations of Mg^2+^ and Na^+^.^58^ Further, specific combinations of metal ions (eg: Ca^2+^ and Na^2+^) promoted the formation of DNA origami monolayers with higher order and at shorter incubation times than other ion combinations.^57^ Our work tested only the effect of individual cations on DNA nanostructure self-assembly, and future work could test a combination of the cations we used here for better assembly yields. Our results for the DNA origami triangle showed no assembly in K^+^ ions, consistent with previous studies.^30^ If needed for certain applications, DNA origami structures can be assembled in Mg^2+^ and then buffer-exchanged into water^59^ or a solution containing K^+.30^

Enhanced biostability is a key feature for DNA nanostructures to be useful *in vivo* so that they withstand assault from the variety of nucleases present in the body. Existing works on biostability enhancement predominantly focus on chemically modifying DNA strands or coating DNA nanostructures with other materials, with a few recent examples focusing on design-based biostability enhancement.^60^ Chemical modification of DNA strands and functionalization of assembled DNA nanostructures may be challenging at times, necessitating design-or assembly-based biostability enhancement. This work provides an assembly-based strategy, showing that structures assembled in monovalent ions can confer high nuclease resistance against DNase I and improved biostability in FBS compared to those assembled in divalent ions. In summary, our study demonstrates successful assembly of a wide variety of DNA nanostructures in different cations, and provides new information on solution-based assembly parameters for improved biostability against nucleases and in biofluids. The cation-dependent assembly conditions tested here could be a useful resource for application-dependent assembly of DNA nanostructures.

## AUTHOR CONTRIBUTIONS

A.R., B.R.M. and H.T. performed non-denaturing PAGE experiments and analysis on the DX, three-point-star and DNA tetrahedron structures. J.M. prepared reagents and performed CD and UV melting experiments. D.G. and T.S. performed agarose gel and AFM analysis of DNA origami structures. J.S., X.W. and A.R.C. acquired funding. X.W. and D.G. wrote the DNA origami content of the manuscript. J.M. and B.R.M. edited the manuscript. X.W. supervised the project, designed experiments, and edited the manuscript. A.R.C. conceived and supervised the project, designed experiments, analyzed and visualized data, and wrote the manuscript.

