## Supporting Information for "Self-assembly of DNA nanostructures in different cations"

### MATERIALS AND METHODS

#### Materials

DNA strands were purchased from Integrated DNA Technologies (IDT) with standard desalting. M13mp18 single stranded circular DNA was purchased from IDT. Sequences of DNA strands used for the DX, three-point-star motif and tetrahedron are provided in **Tables S2** and **S3**. Sequences of DNA strands for the origami triangle are provided in **Table S4**. For different metal ions, the chemicals used were cadmium (II) nitrate tetrahydrate (Cd $(NO_3)_2 \cdot 4H_2O$), ≥98% (Fisher Scientific), calcium chloride dihydrate ($CaCl_2 \cdot 2H_2O$), ≥99% (Sigma-Aldrich), lithium chloride (LiCl), ≥99% (Sigma Aldrich), silver nitrate, ($AgNO_3$), (Millipore), nickel chloride hexahydrate, ($NiCl_2 \cdot 6H_2O$) (VWR, high purity), copper (II) sulfate pentahydrate ($CuSO_4 \cdot 5H_2O$), ≥98% (Sigma-Aldrich), barium chloride ($BaCl_2$), 99% (Sigma-Aldrich), lead (II) chloride ($PbCl_2$), 98% (Sigma-Aldrich), sodium chloride (NaCl), ≥99% (Sigma-Aldrich), zinc chloride anhydrous ($ZnCl_2$), 97% (Sterm chemicals), magnesium chloride ($MgCl_2$), ≥99% (Sigma-Aldrich), and potassium chloride (KCl), 99-100.5% (Sigma-Aldrich).

#### DNA nanostructure assembly

***Double crossover (DX) motif.*** Component DNA strands (DX1, DX2, DX3 and DX4) were mixed in equimolar ratios in 1× Tris-Acetic-EDTA (TAE) buffer (40 mM Tris base, pH 8.0; 20 mM acetic acid, 2 mM EDTA) containing 10 mM of different cations ($Mg^{2+}$, $Ca^{2+}$, $Ba^{2+}$, $Ni^{2+}$, $Cd^{2+}$, $Pb^{2+}$, $Zn^{2+}$, $Cu^{2+}$, $Na^{+}$, $K^{+}$, $Li^{+}$ and $Ag^{+}$). The final concentration of the DNA complex was 250 nM for polyacrylamide gel electrophoresis (PAGE) studies, 1 μM for UV melting studies and 5 μM for circular dichroism (CD) studies. The mixture was annealed from 90 °C to 20 °C over two hours in a T100 Thermal Cycler (Bio-Rad) using the following protocol: 90 °C for 5 min, 65 °C for 20 min, 45 °C for 20 min, 37 °C for 30 min, and 20 °C for 30 min.

***3-point-star motif.*** Component DNA strands (L, M and S-blunt) were mixed in 1:3:3 ratio in 1× TAE containing 10 mM of different cations. Sequences of DNA strands are given in **Table S2**. The final concentration of the DNA complex was 250 nM. The mixture was annealed from 90 °C to 20 °C over two hours in a T100 Thermal Cycler (Bio-Rad) using the following protocol: 90 °C for 5 min, 65 °C for 20 min, 45 °C for 20 min, 37 °C for 30 min, and 20 °C for 30 min.

***DNA tetrahedron.*** DNA strands L, M and S were mixed in 1:3:3 ratio at 30 nM in 1× TAE containing 10 mM of different cations. Sequences of DNA strands are given in **Table S3**. The DNA solution was slowly cooled down from 95°C to room temperature over 48 hours by placing the tubes in 2 liters of hot water in a beaker placed in a Styrofoam box.

***DNA origami.*** M13mp18 scaffold strand and 208 staple strands were mixed in a 1:5 ratio (final DNA concentration of 5 nM) in 1× TAE buffer (40 mM Tris, 20 mM acetic acid, 1 mM EDTA) containing 10 mM or higher concentrations of a cation. The mixture was annealed from 80°C to room temperature over 16 hours in a Biometra TRIO Thermal Cycler.

**Polyacrylamide gel electrophoresis (PAGE)**

Non-denaturing gels containing 4-10% polyacrylamide (19:1 acrylamide/bisacrylamide) were run at 4 °C (100 V, constant voltage) in 1× TAE running buffer with or without 10 mM $Mg^{2+}$ (see text for details). Samples were mixed with gel loading dye containing 50% glycerol, bromophenol blue and 1× TAE before loading on gels. Gels were stained in deionized water containing 0.5× GelRed (Biotium). Gels were imaged on a Bio-Rad Gel Doc XR+ imager using the default settings for GelRed with UV illumination. Gel images were exported as 12-bit images and quantified using ImageJ. Quantification was done using the highest exposure image that did not contain saturated pixels in the band of interest. For the DX assembly, we quantified the bands corresponding to the structure. For three-point-star assembly, we calculated the percentage of the band corresponding to the structure in the entire lane.

**Agarose gel electrophoresis**

The DNA origami triangles were run on 1% agarose gels in 1× TAE running buffer. The gels were pre-stained with SYBR-Green (Thermo Fisher), run at a constant voltage of 7.5 V/cm for 2 hours on ice and scanned using GelDoc (UVP GelStudio).

**UV melting experiments**

Experiments were performed in a Cary 3500 UV-Visible Spectrophotometer equipped with a temperature controller, using 1 μM DNA complexes assembled in 1× TAE buffer containing 10 mM metal ions. Melting curves were acquired at 260 nm by heating and cooling from 20 °C to 90 °C at a rate of 1 °C/min.

**Circular dichroism experiments**

CD spectra were collected on 5 μM DNA complexes assembled in 1× TAE buffer containing 10 mM metal ions. Experiments were performed on a Jasco-815 CD spectrometer at room temperature in a quartz cell with a 1 mm path length. CD spectra were collected from 350 to 200 nm with a scanning speed of 100 nm/min and with 3 accumulations. The bandwidth was 1.0 nm, and the digital integration time was 1.0 s. All CD spectra were baseline-corrected for signal contributions due to the buffer.

#### AFM imaging

The mica surface was washed with double distilled water and dried by compressed air. 5 µl of the annealed DNA sample (5 nM) was deposited onto the freshly-cleaved mica surface and incubated for 5 minutes. 40 µl of 1× TAE buffer with different cations (specific to the condition the DNA origami was annealed in) was then added and the AFM scan was performed under tapping mode in the fluid cell on Multimode AFM Asylum Research Cypher S with SNL-10 probe (Bruker Nano, Inc.). After the scan, raw AFM files were processed with Gwyddion V.2.6.1 software by using the flatten function to adjust the AFM images to reflect the actual height of the DNA origami nanostructures.

#### Nuclease degradation experiments

Annealed DX motif (at 0.5 µM) were first mixed with DNase I reaction buffer (final of 1×). DNase I enzyme was purchased from New England Biolabs (Catalog # M0303S), and according to the vendor, is isolated from a recombinant *E. Coli* strain carrying an MBP fusion clone of bovine pancreatic DNase I. Dilutions of DNase I enzyme to different units was made in nuclease-free water. For the DNase I assay, 1 µl of the enzyme was added to 10 µl of the sample containing DNase I reaction buffer. Samples were incubated at 37 °C for different time intervals. Incubated samples were mixed with gel loading dye and run on non-denaturing polyacrylamide gels to analyze degradation over time. A step-by-step protocol for our gel-based method of DNA nanostructure degradation analysis is provided in *Current Protocols in Nucleic Acid Chemistry*, 82: e115 (2020).

#### Stability test in FBS

Fetal bovine serum (FBS) was purchased from Thermo Fisher (Gibco, cat #: A4736301). FBS was added to annealed DNA complexes (DX motif or DNA origami triangle) to be at a final concentration of 10%. Typically, we added 1µl of FBS to 9 µl of the annealed samples and incubated them at 37 °C for different time intervals. Incubated samples were mixed with gel loading dye and run on non-denaturing polyacrylamide (for DX) or agarose gels (for DNA origami) to analyze degradation over time. Data was quantified using a minimum of two replicates for each condition tested.

### STATISTICAL ANALYSIS

All experiments discussed in the manuscript were performed with at least two replicates and data are presented as mean and standard deviation of error propagated from replicates. Statistical details of experiments can be found in figure legends where applicable.

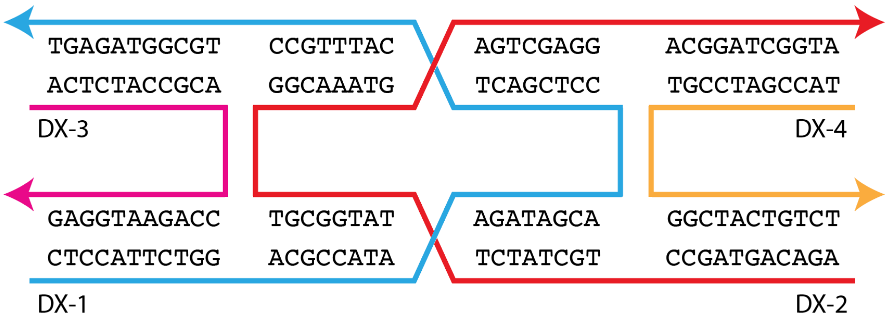


**Figure S1**. Schematic and sequence of the DX motif.

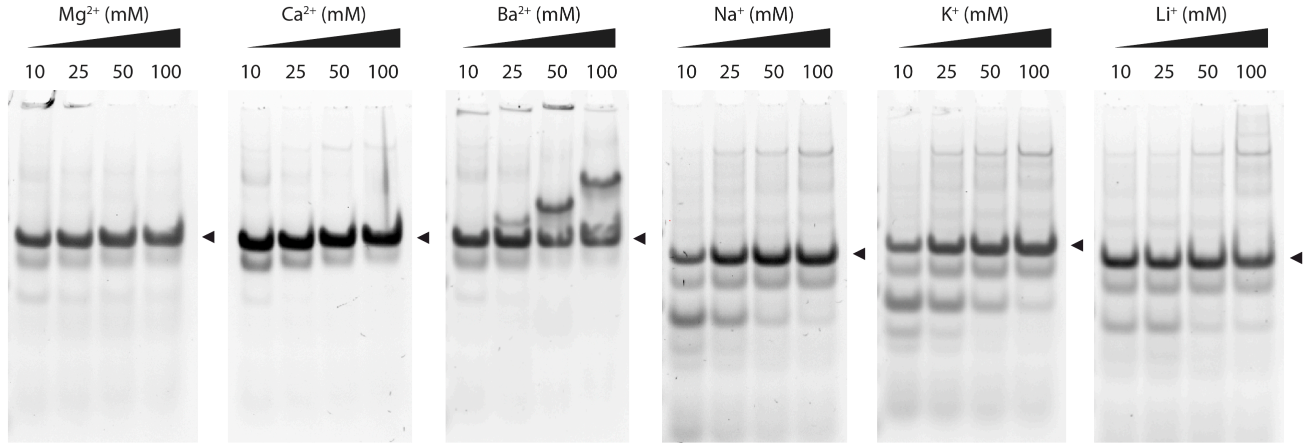


**Figure S2**. Assembly of DX in different metal ions. Full gels of images shown in Figure 1f.

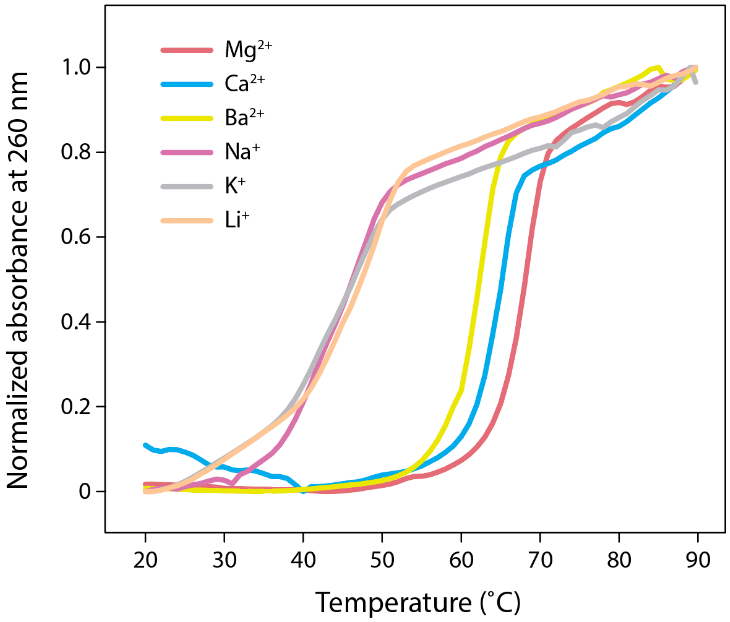


**Figure S3**. UV melting profiles of DX assembled in buffer containing different metal ions.

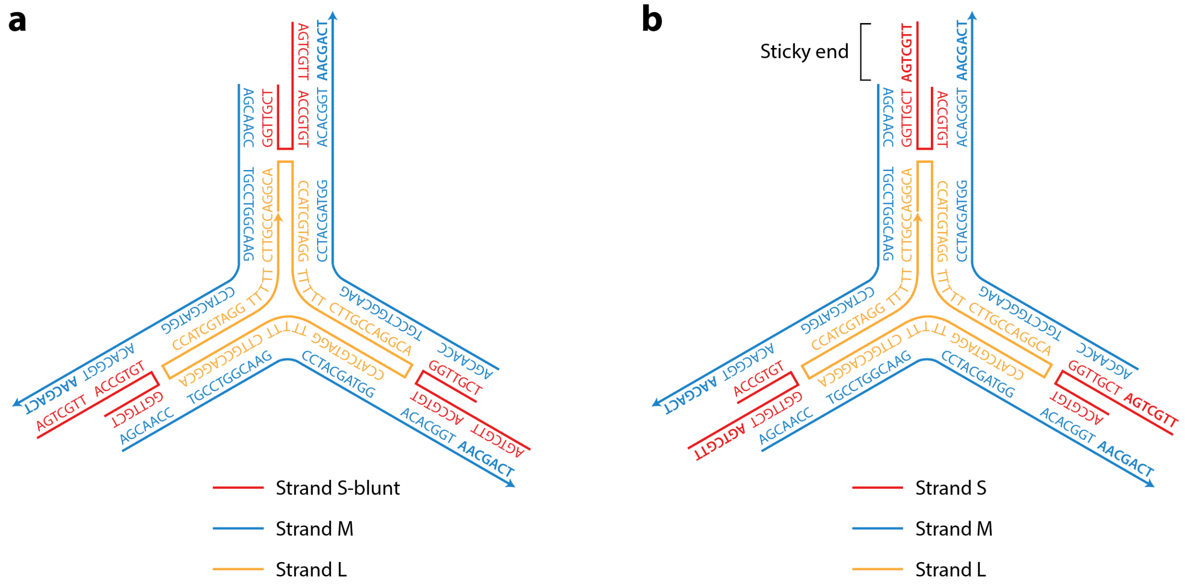


**Figure S4**. (a) Design of the 3-point star motif without sticky ends to assemble the motif. (b) 3-point-star motif with sticky ends that allows hierarchical assembly into a DNA tetrahedron.

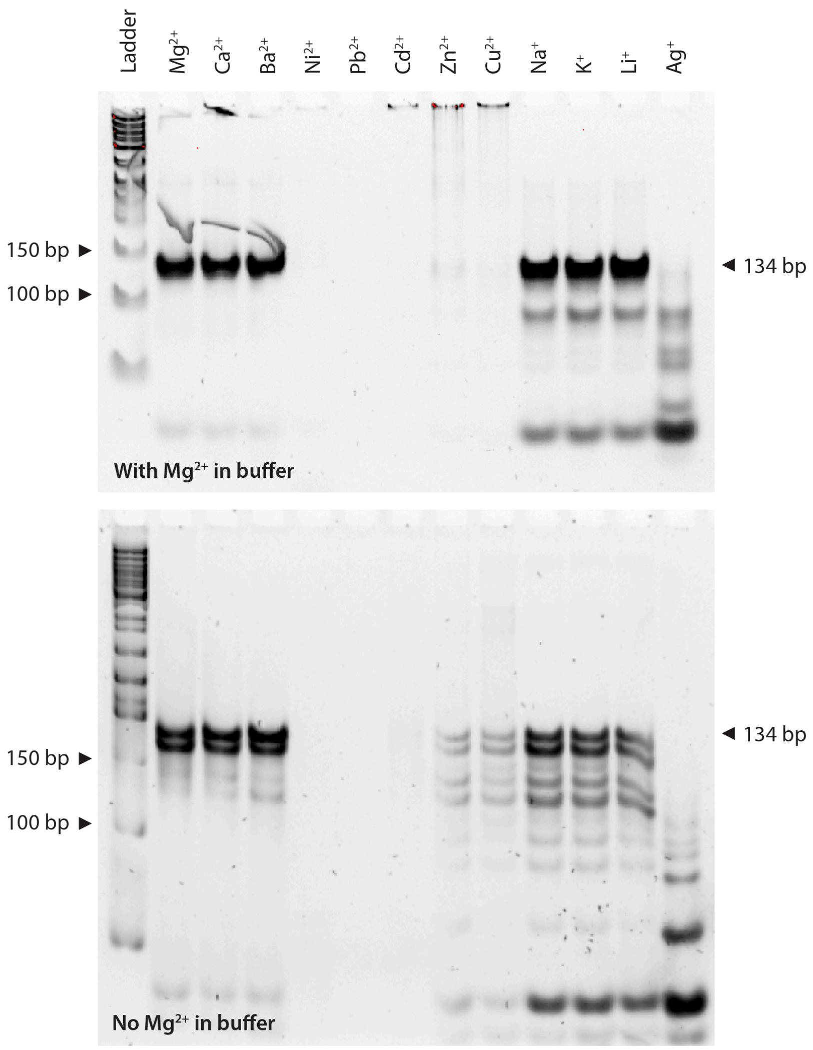


**Figure S5**. Three-point star motif assembled in different ions and run in gels with or without $Mg^{2+}$.

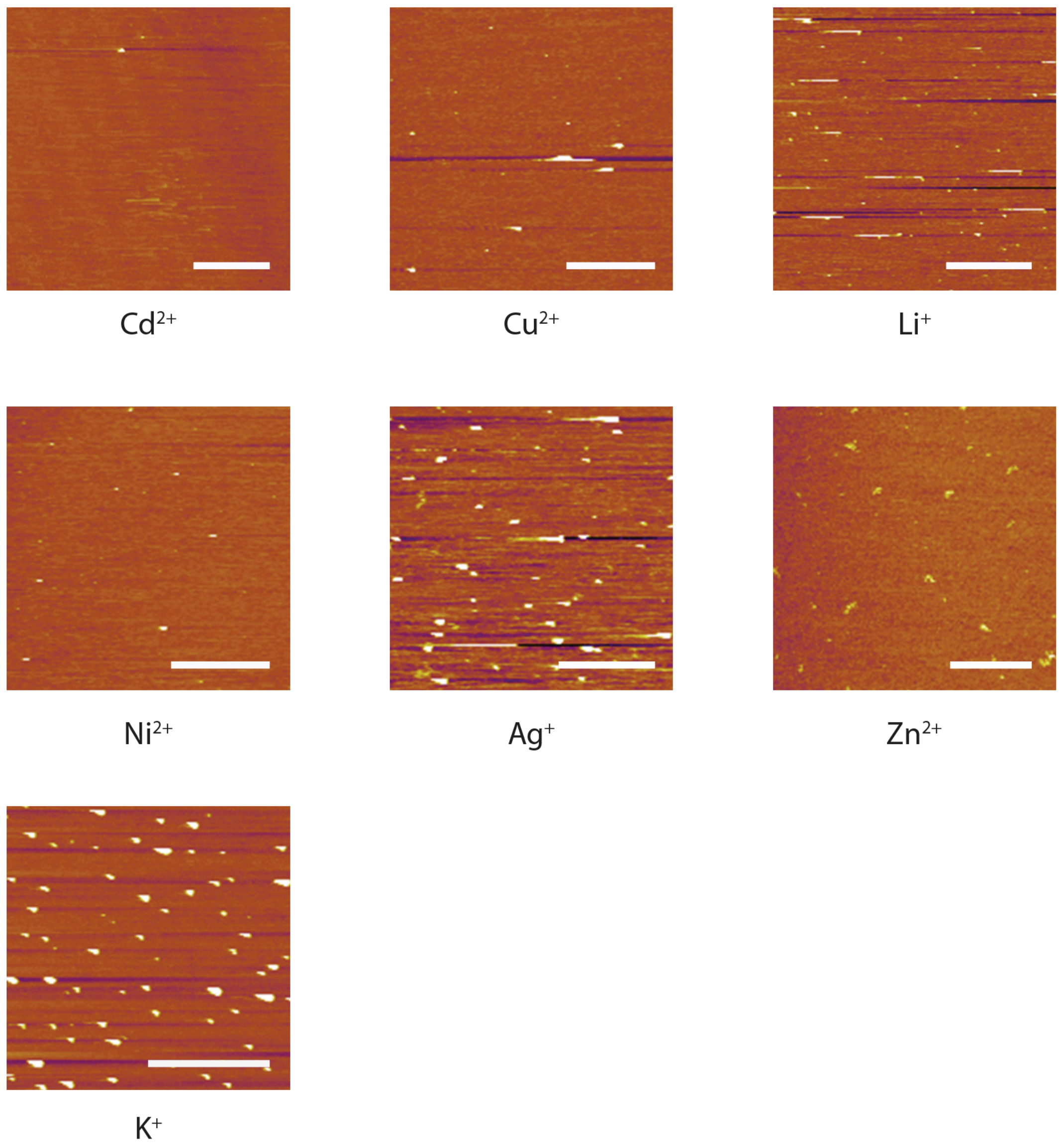


**Figure S6**. AFM analysis of DNA origami triangles assembled in 10 mM of different ions. Scale bars in insets are 100 nm.

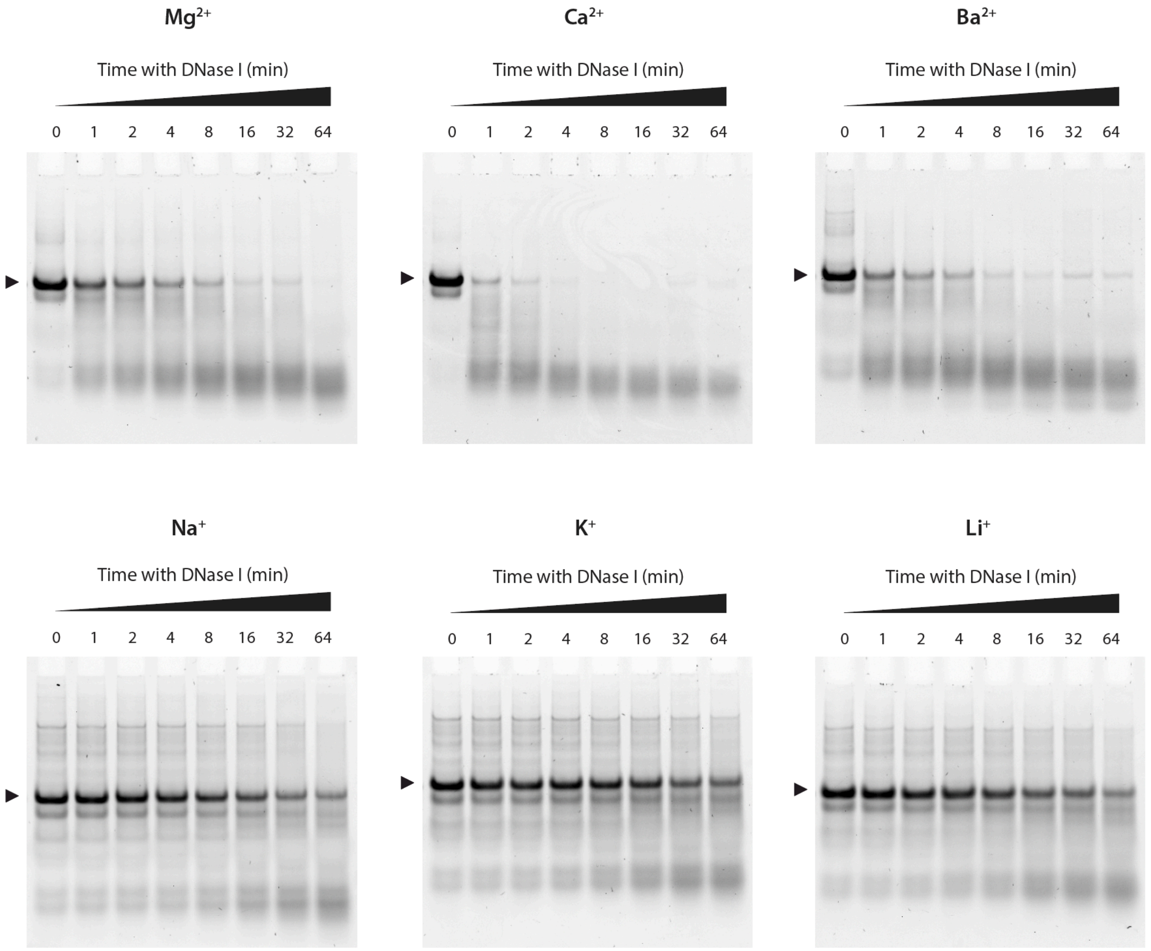


**Figure S7**. Nuclease degradation analysis of DX motif assembled in different ions. Full gels of images shown in Figure 4b.

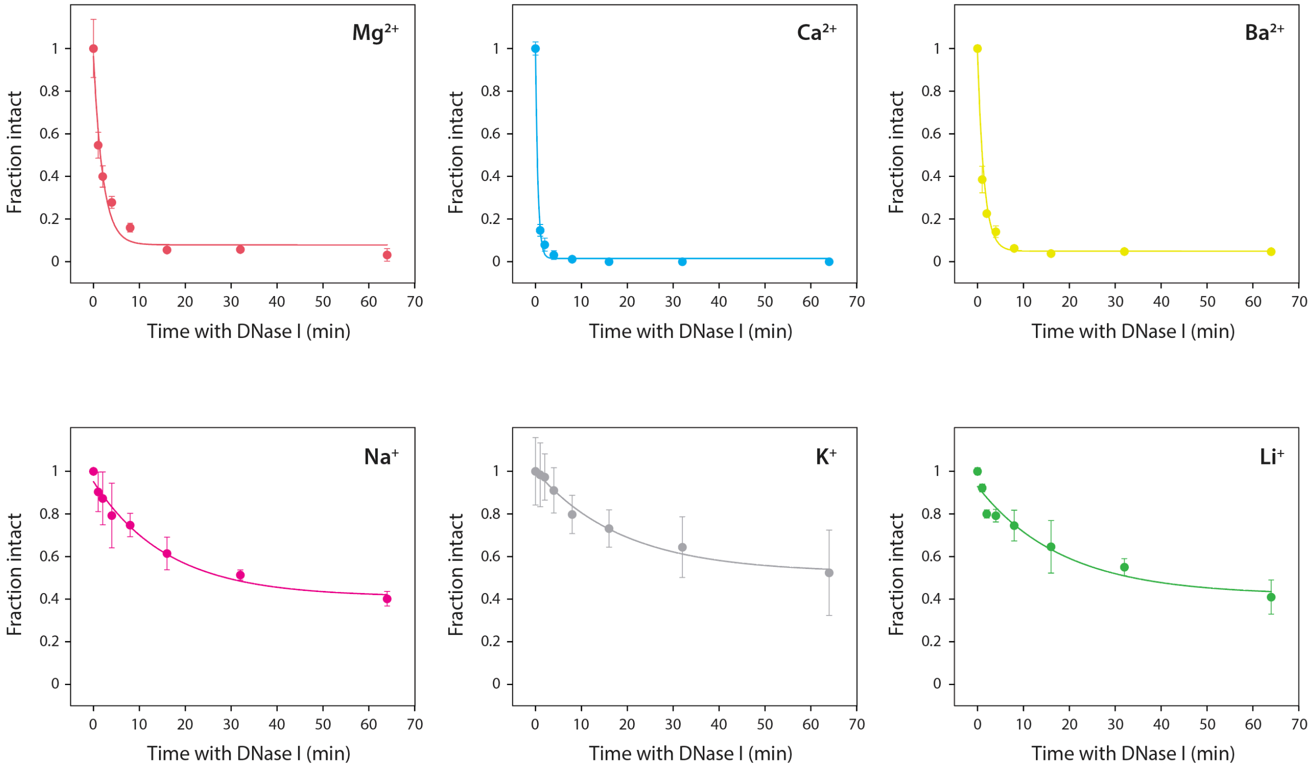


**Figure S8**. Quantitative analysis of nuclease degradation of DX motif assembled in different ions.

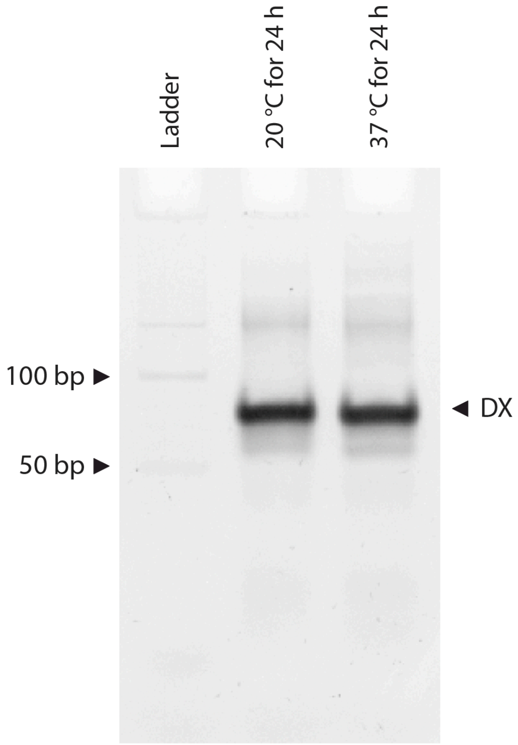


**Figure S9.** Stability of DX motif at 37 °C for 24 h.

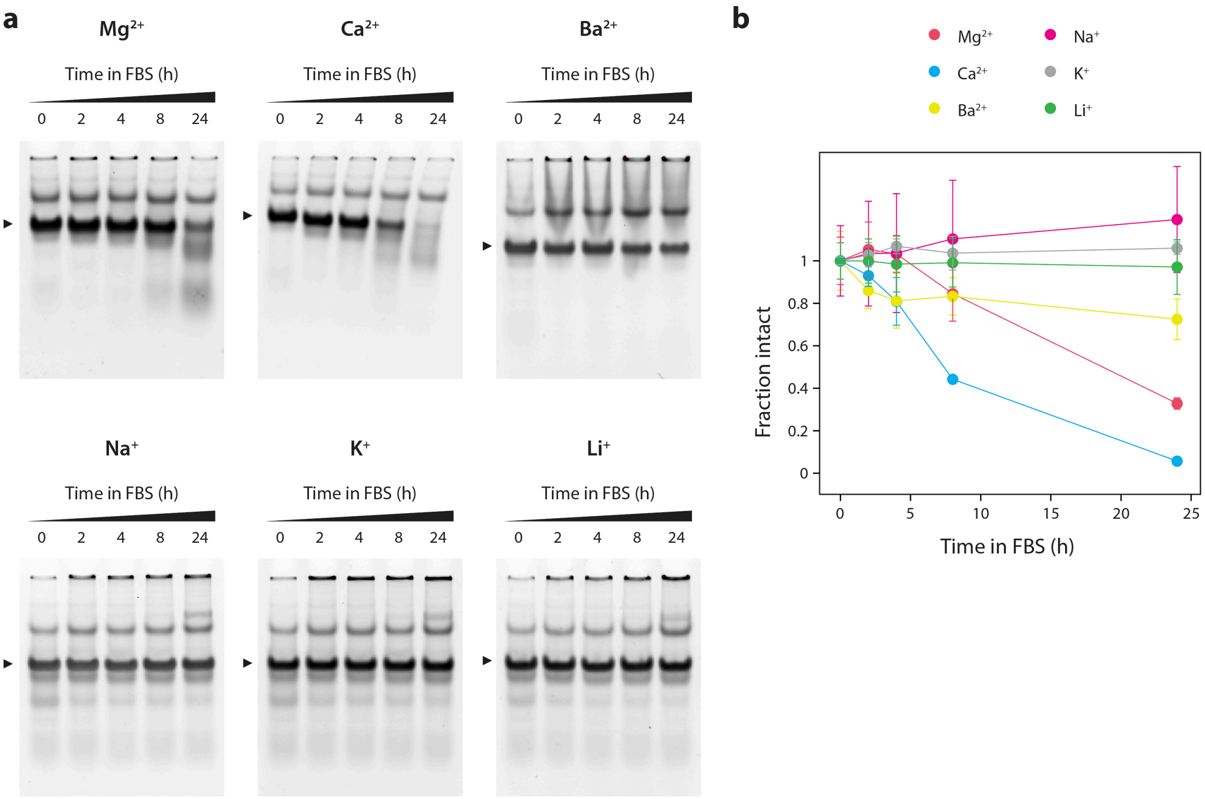


**Figure S10**. DX biostability in FBS. (a) PAGE analysis of DX degradation in FBS. Full gels of images shown in Figure 4e. (b) Quantified results from gels shown in (a). Fraction intact at the 24-hour time point is shown in Figure 4f. Components in FBS also get stained in the gel and appear as a higher mobility band.

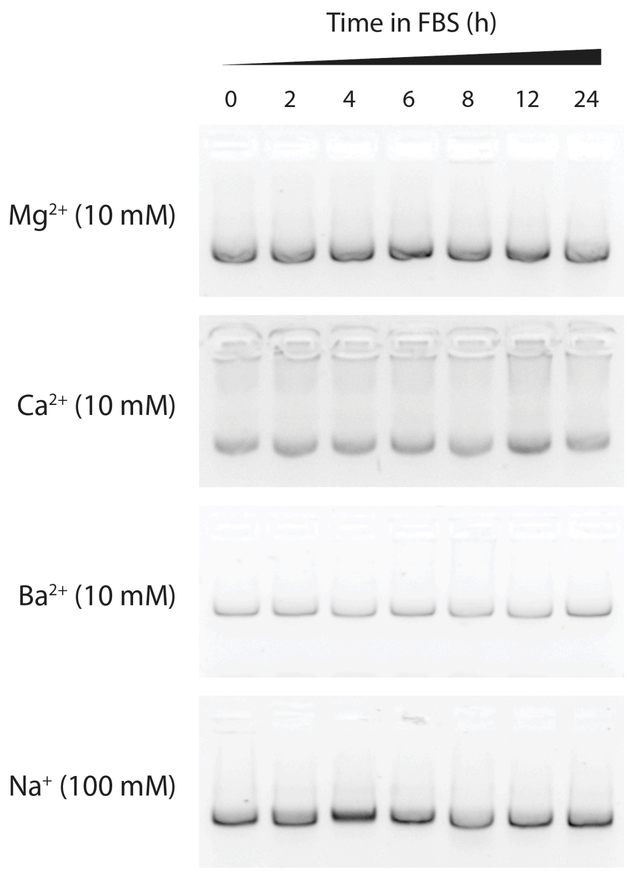


**Figure S11**. Agarose gel analysis of DNA origami triangle degradation in 10% FBS.

**Table S1**. Time constants for nuclease degradation analysis of DX motif in different ions.

| Metal ion | Time constant ( $\tau$ ) | R square |
|---|---|---|
| $Mg^{2+}$ | $2.03 \pm 0.35$ | 0.9639 |
| $Ca^{2+}$ | $0.52 \pm 0.05$ | 0.9950 |
| $Ba^{2+}$ | $1.41 \pm 0.28$ | 0.9480 |
| $Na^{+}$ | $16.04 \pm 3.20$ | 0.9731 |
| $K^{+}$ | $18.85 \pm 3.48$ | 0.9791 |
| $Li^{+}$ | $18.42 \pm 6.82$ | 0.9183 |

**Table S2**. Sequences of DNA strands used for DX motif.

| Strand | Sequence | Length |
|---|---|---|
| DX1 | CTCCATTCTGGACGCCATAAGATAGCACCTCGACTCATTTGCCTGCGGTAGAGT | 54 |
| DX2 | AGACAGTAGCCTGCTATCTTATGGCGTGGCAAATGAGTCGAGGACGGATCGGTA | 54 |
| DX3 | ACTCTACCGCACCAGAATGGAG | 22 |
| DX4 | TACCGATCCGTGGCTACTGTCT | 22 |

**Table S3**. Sequences of DNA strands used for three-point-star motif (strands L, M and S-blunt) and the DNA tetrahedron (strands L, M and S).

| Strand | Sequence | Length |
|---|---|---|
| Strand L | AGGCACCATCGTAGGTTTTTCTTGCCAGGCACCATCGTA GGTTTTTCTTGCCAGGCACCATCGTAGGTTTTTCTTGCC | 78 |
| Strand M | AGCAACCTGCCTGGCAAGCCTACGATGGACACGGTAACGACT | 42 |
| Strand S | ACCGTGTGGTTGCTAGTCGTT | 21 |
| Strand S-blunt | AGTCGTTACCGTGTGGTTGCT | 21 |

**Table S4**. Staple DNA oligos used for self-assembly of DNA triangle origami.

| Strand | Sequence |
|---|---|
| A_01 | CGGGGTTTCCTCAAGAGAAGGATTTTGAATTA |
| A_02 | AGCGTCATGTCTCTGAATTTACCGACTACCTT |
| A_03 | TTCATAATCCCCTTATTAGCGTTTTTCTTACC |
| A_04 | ATGGTTTATGTCACAATCAATAGATATTAAAC |
| A_05 | TTTGATGATTAAGAGGCTGAGACTTGCTCAGTACCAGGCG |
| A_06 | CCGGAACCCAGAATGGAAAGCGCAACATGGCT |
| A_07 | AAAGACAACATTTTCGGTCATAGCCAAAATCA |
| A_08 | GACGGGAGAATTAACTCGGAATAAGTTTATTTCCAGCGCC |
| A_09 | GATAAGTGCCGTCGAGCTGAAACATGAAAGTATACAGGAG |
| A_10 | TGTACTGGAAATCCTCATTAAAGCAGAGCCAC |
| A_11 | CACCGGAAAGCGCGTTTTCATCGGAAGGGCGA |
| A_12 | CATTCAACAAACGCAAAGACACCAGAACACCCTGAACAAA |
| A_13 | TTTAACGGTTCGGAACCTATTATTAGGGTTGATATAAGTA |
| A_14 | CTCAGAGCATATTCACAAACAAATTAATAAGT |
| A_15 | GGAGGGAATTTAGCGTCAGACTGTCCGCCTCC |
| A_16 | GTCAGAGGGTAATTGATGGCAACATATAAAAGCGATTGAG |
| A_17 | TAGCCCGGAATAGGTGAATGCCCCCTGCCTATGGTCAGTG |
| A_18 | CCTTGAGTCAGACGATTGGCCTTGCGCCACCC |
| A_19 | TCAGAACCCAGAATCAAGTTTGCCGGTAAATA |
| A_20 | TTGACGGAAATACATACATAAAGGGCGCTAATATCAGAGA |
| A_21 | CAGAGCCAGGAGGTTGAGGCAGGTAACAGTGCCCG |
| A_22 | ATTAAAGGCCGTAATCAGTAGCGAGCCACCCT |
| A_23 | GATAACCCACAAGAATGTTAGCAAACGTAGAAAATTATTC |
| A_24 | GCCGCCAGCATTGACACCACCCTC |
| A_25 | AGAGCCGCACCATCGATAGCAGCATGAATTAT |
| A_26 | CACCGTCACCTTATTACGCAGTATTGAGTTAAGCCCAATA |
| A_27 | AGCCATTTAAACGTCACCAATGAACACCAGAACCA |
| A_28 | ATAAGAGCAAGAAACATGGCATGATTAAGACTCCGACTTG |
| A_29 | CCATTAGCAAGGCCGGGGGAATTA |
| A_30 | GAGCCAGCGAATACCCAAAAGAACATGAAATAGCAATAGC |
| A_31 | TATCTTACCGAAGCCCAAACGCAATAATAACGAAAATCACCAG |
| A_32 | CAGAAGGAAACCGAGGTTTTTAAGAAAAGTAAGCAGATAGCCG |
| A_33 | CCTTTTTTCATTTAACAATTTCATAGGATTAG |
| A_34 | TTTAACCTATCATAGGTCTGAGAGTTCCAGTA |
| A_35 | AGTATAAAATATGCGTTATACAAAGCCATCTT |
| A_36 | CAAGTACCTCATTCCAAGAACGGGAAATTCAT |
| A_37 | AGAGAATAACATAAAAACAGGGAAGCGCATTA |
| A_38 | AAAACAAAATTAATTAAATGGAAACAGTACATTAGTGAAT |
| A_39 | TTATCAAACCGGCTTAGGTTGGGTAAGCCTGT |
| A_40 | TTAGTATCGCCAACGCTCAACAGTCGGCTGTC |

| | |
|---|---|
| A_41 | TTTCCTTAGCACTCATCGAGAACAATAGCAGCCTTTACAG |
| A_42 | AGAGTCAAAAATCAATATATGTGATGAAACAAACATCAAG |
| A_43 | ACTAGAAATATATAACTATATGTACGCTGAGA |
| A_44 | TCAATAATAGGGCTTAATTGAGAATCATAATT |
| A_45 | AACGTCAAAAATGAAAAGCAAGCCGTTTTTATGAAACCAA |
| A_46 | GAGCAAAAGAAGATGAGTGAATAACCTTGCTTATAGCTTA |
| A_47 | GATTAAGAAATGCTGATGCAAATCAGAATAAA |
| A_48 | CACCGGAATCGCCATATTTAACAAAATTTACG |
| A_49 | AGCATGTATTTCATCGTAGGAATCAAACGATTTTTTGTTT |
| A_50 | ACATAGCGCTGTAAATCGTCGCTATTCATTTCAATTACCT |
| A_51 | GTTAAATACAATCGCAAGACAAAGCCTTGAAA |
| A_52 | CCCATCCTCGCCAACATGTAATTTAATAAGGC |
| A_53 | TCCCAATCCAAATAAGATTACCGCGCCCAATAAATAATAT |
| A_54 | TCCCTTAGAATAACGCGAGAAAACTTTTACCGACC |
| A_55 | GTGTGATAAGGCAGAGGCATTTTCAGTCCTGA |
| A_56 | ACAAGAAAGCAAGCAAATCAGATAACAGCCATATTATTTA |
| A_57 | GTTTGAAATTCAAATATATTTTAG |
| A_58 | AATAGATAGAGCCAGTAATAAGAGATTTAATG |
| A_59 | GCCAGTTACAAAATAATAGAAGGCTTATCCGGTTATCAAC |
| A_60 | TTCTGACCTAAAATATAAAGTACCGACTGCAGAAC |
| A_61 | GCGCCTGTTATTCTAAGAACGCGATTCCAGAGCCTAATTT |
| A_62 | TCAGCTAAAAAAGGTAAAGTAATT |
| A_63 | ACGCTAACGAGCGTCTGGCGTTTTAGCGAACCCAACATGT |
| A_64 | ACGACAATAAATCCCGACTTGCGGGAGATCCTGAATCTTACCA |
| A_65 | TGCTATTTTGCACCCAGCTACAATTTTGTTTTGAAGCCTTAAA |
| B_01 | TCATATGTGTAATCGTAAAACTAGTCATTTTC |
| B_02 | GTGAGAAAATGTGTAGGTAAAGATACAACTTT |
| B_03 | GGCATCAAATTTGGGGCGCGAGCTAGTTAAAG |
| B_04 | TTCGAGCTAAGACTTCAAATATCGGGAACGAG |
| B_05 | ACAGTCAAAGAGAATCGATGAACGACCCCGGTTGATAATC |
| B_06 | ATAGTAGTATGCAATGCCTGAGTAGGCCGGAG |
| B_07 | AACCAGACGTTTAGCTATATTTTCTTCTACTA |
| B_08 | GAATACCACATTCAACTTAAGAGGAAGCCCGATCAAAGCG |
| B_09 | AGAAAAGCCCCAAAAAGAGTCTGGAGCAAACAATCACCAT |
| B_10 | CAATATGACCCTCATATATTTTAAAGCATTAA |
| B_11 | CATCCAATAAATGGTCAATAACCTCGGAAGCA |
| B_12 | AACTCCAAGATTGCATCAAAAAGATAATGCAGATACATAA |
| B_13 | CGTTCTAGTCAGGTCATTGCCTGACAGGAAGATTGTATAA |
| B_14 | CAGGCAAGATAAAAATTTTTAGAATATTCAAC |
| B_15 | GATTAGAGATTAGATACATTTCGCAAATCATA |
| B_16 | CGCCAAAAGGAATTACAGTCAGAAGCAAAGCGCAGGTCAG |
| B_17 | GCAAATATTTAAATTGAGATCTACAAAGGCTACTGATAAA |
| B_18 | TTAATGCCTTATTTCAACGCAAGGGCAAAGAA |

| | |
|---|---|
| B_19 | TTAGCAAATAGATTTAGTTTGACCAGTACCTT |
| B_20 | TAATTGCTTTACCCTGACTATTATGAGGCATAGTAAGAGC |
| B_21 | ATAAAGCCTTTGCGGGAGAAGCCTGGAGAGGGTAG |
| B_22 | TAAGAGGTCAATTCTGCGAACGAGATTAAGCA |
| B_23 | AACACTATCATAACCCATCAAAAATCAGGTCTCCTTTTGA |
| B_24 | ATGACCCTGTAATACTTCAGAGCA |
| B_25 | TAAAGCTATATAACAGTTGATTCCCATTTTTG |
| B_26 | CGGATGGCACGAGAATGACCATAATCGTTTACCAGACGAC |
| B_27 | TAATTGCTTGGAAGTTTCATTCCAAATCGGTTGTA |
| B_28 | GATAAAAACCAAAATATTAAACAGTTCAGAAATTAGAGCT |
| B_29 | ACTAAAGTACGGTGTCGAATATAA |
| B_30 | TGCTGTAGATCCCCCTCAAATGCTGCGAGAGGCTTTTGCA |
| B_31 | AAAGAAGTTTTGCCAGCATAAATATTCATTGACTCAACATGTT |
| B_32 | AATACTGCGGAATCGTAGGGGGTAATAGTAAAATGTTTAGACT |
| B_33 | AGGGATAGCTCAGAGCCACCACCCCATGTCAA |
| B_34 | CAACAGTTTATGGGATTTTGCTAATCAAAAGG |
| B_35 | GCCGCTTTGCTGAGGCTTGCAGGGGAAAAGGT |
| B_36 | GCGCAGACTCCATGTTACTTAGCCCGTTTTAA |
| B_37 | ACAGGTAGAAAGATTCATCAGTTGAGATTTAG |
| B_38 | CCTCAGAACCGCCACCCAAGCCCAATAGGAACGTAAATGA |
| B_39 | ATTTTCTGTCAGCGGAGTGAGAATACCGATAT |
| B_40 | ATTCGGTCTGCGGGATCGTCACCCGAAATCCG |
| B_41 | CGACCTGCGGTCAATCATAAGGGAACGGAACAACATTATT |
| B_42 | AGACGTTACCATGTACCGTAACACCCCTCAGAACCGCCAC |
| B_43 | CACGCATAAGAAAGGAACAACTAAGTCTTTCC |
| B_44 | ATTGTGTCTCAGCAGCGAAAGACACCATCGCC |
| B_45 | TTAATAAAACGAACTAACCGAACTGACCAACTCCTGATAA |
| B_46 | AGGTTTAGTACCGCCATGAGTTTCGTCACCAGGATCTAAA |
| B_47 | GTTTTGTCAGGAATTGCGAATAATCCGACAAT |
| B_48 | GACAACAAGCATCGGAACGAGGGTGAGATTTG |
| B_49 | TATCATCGTTGAAAGAGGACAGATGGAAGAAAAATCTACG |
| B_50 | AGCGTAACTACAAACTACAACGCCTATCACCGTACTCAGG |
| B_51 | TAGTTGCGAATTTTTTCACGTTGATCATAGTT |
| B_52 | GTACAACGAGCAACGGCTACAGAGGATACCGA |
| B_53 | ACCAGTCAGGACGTTGGAACGGTGTACAGACCGAAACAAA |
| B_54 | ACAGACAGCCCAAATCTCCAAAAAAAAATTTCTTA |
| B_55 | AACAGCTTGCTTTGAGGACTAAAGCGATTATA |
| B_56 | CCAAGCGCAGGCGCATAGGCTGGCAGAACTGGCTCATTAT |
| B_57 | CGAGGTGAGGCTCCAAAAGGAGCC |
| B_58 | ACCCCCAGACTTTTTCATGAGGAACTTGCTTT |
| B_59 | ACCTTATGCGATTTTATGACCTTCATCAAGAGCATCTTTG |
| B_60 | CGGTTTATCAGGTTTCCATTAAACGGGAATACACT |
| B_61 | AAAACACTTAATCTTGACAAGAACTTAATCATTGTGAATT |

| | |
|---|---|
| B_62 | GGCAAAAGTAAAATACGTAATGCC |
| B_63 | TGGTTTAATTTCAACTCGGATATTCATTACCCACGAAAGA |
| B_64 | ACCAACCTAAAAAATCAACGTAACAAATAAATTGGGCTTGAGA |
| B_65 | CCTGACGAGAAACACCAGAACGAGTAGGCTGCTCATTCAGTGA |
| Link-A1C | TTAATTAATTTTTTACCATATCAAA |
| Link-A2C | TTAATTTCATCTTAGACTTTACAA |
| Link-A3C | CTGTCCAGACGTATACCGAACGA |
| Link-A4C | TCAAGATTAGTGTAGCAATACT |
| Link-B1A | TGTAGCATTCCTTTTATAAACAGTT |
| Link-B2A | TTTAATTGTATTTCCACCAGAGCC |
| Link-B3A | ACTACGAAGGCTTAGCACCATTA |
| Link-B4A | ATAAGGCTTGCAACAAAGTTAC |
| Link-C1B | GTGGGAACAAATTTCTATTTTTGAG |
| Link-C2B | CGGTGCGGGCCTTCCAAAAACATT |
| Link-C3B | ATGAGTGAGCTTTTAAATATGCA |
| Link-C4B | ACTATTAAAGAGGATAGCGTCC |
| Loop | GCGCTTAATGCGCCGCTACAGGGC |
| C_01 | TCGGGAGATATACAGTAACAGTACAAATAATT |
| C_02 | CCTGATTAAAGGAGCGGAATTATCTCGGCCTC |
| C_03 | GCAAATCACCTCAATCAATATCTGCAGGTCGA |
| C_04 | CGACCAGTACATTGGCAGATTCACCTGATTGC |
| C_05 | TGGCAATTTTTAACGTCAGATGAAAACAATAACGGATTCG |
| C_06 | AAGGAATTACAAAGAAACCACCAGTCAGATGA |
| C_07 | GGACATTCACCTCAAATATCAAACACAGTTGA |
| C_08 | TTGACGAGCACGTATACTGAAATGGATTATTTAATAAAAG |
| C_09 | CCTGATTGCTTTGAATTGCGTAGATTTTCAGGCATCAATA |
| C_10 | TAATCCTGATTATCATTTTGCGGAGAGGAAGG |
| C_11 | TTATCTAAAGCATCACCTTGCTGATGGCCAAC |
| C_12 | AGAGATAGTTTGACGCTCAATCGTACGTGCTTTCCTCGTT |
| C_13 | GATTATACACAGAAATAAAGAAATACCAAGTTACAAAATC |
| C_14 | TAGGAGCATAAAAGTTTGAGTAACATTGTTTG |
| C_15 | TGACCTGACAAATGAAAAATCTAAAATATCTT |
| C_16 | AGAATCAGAGCGGGAGATGGAAATACCTACATAACCCTTC |
| C_17 | GCGCAGAGGCGAATTAATTATTTGCACGTAAATTCTGAAT |
| C_18 | AATGGAAGCGAACGTTATTAATTTCTAACAAC |
| C_19 | TAATAGATCGCTGAGAGCCAGCAGAAGCGTAA |
| C_20 | GAATACGTAACAGGAAAAACGCTCCTAAACAGGAGGCCGA |
| C_21 | TCAATAGATATTAAATCCTTTGCCGGTTAGAACCT |
| C_22 | CAATATTTGCCTGCAACAGTGCCATAGAGCCG |
| C_23 | TTAAAGGGATTTTAGATACCGCCAGCCATTGCGGCACAGA |
| C_24 | ACAATTCGACAACTCGTAATACAT |
| C_25 | TTGAGGATGGTCAGTATTAACACCTTGAATGG |
| C_26 | CTATTAGTATATCCAGAACAATATCAGGAACGGTACGCCA |

| | |
|---|---|
| C_27 | CGCGAACTAAAACAGAGGTGAGGCTTAGAAGTATT |
| C_28 | GAATCCTGAGAAGTGTATCGGCCTTGCTGGTACTTTAATG |
| C_29 | ACCACCAGCAGAAGATGATAGCCC |
| C_30 | TAAAACATTAGAAGAACTCAAACTTTTTATAATCAGTGAG |
| C_31 | GCCACCGAGTAAAAGAACATCACTTGCCTGAGCGCCATTAAAA |
| C_32 | TCTTTGATTAGTAATAGTCTGTCCATCACGCAAATTAACCGTT |
| C_33 | CGCGTCTGATAGGAACGCCATCAACTTTTACA |
| C_34 | AGGAAGATGGGGACGACGACAGTAATCATATT |
| C_35 | CTCTAGAGCAAGCTTGCATGCCTGGTCAGTTG |
| C_36 | CCTTCACCGTGAGACGGGCAACAGCAGTCACA |
| C_37 | CGAGAAAGGAAGGGAAGCGTACTATGGTTGCT |
| C_38 | GCTCATTTTTTAACCAGCCTTCCTGTAGCCAGGCATCTGC |
| C_39 | CAGTTTGACGCACTCCAGCCAGCTAAACGACG |
| C_40 | GCCAGTGCGATCCCCGGGTACCGAGTTTTTCT |
| C_41 | TTTCACCAGCCTGGCCCTGAGAGAAAGCCGGCGAACGTGG |
| C_42 | GTAACCGTCTTTCATCAACATTAAAATTTTTGTTAAATCA |
| C_43 | ACGTTGTATTCCGGCACCGCTTCTGGCGCATC |
| C_44 | CCAGGGTGGCTCGAATTCGTAATCCAGTCACG |
| C_45 | TAGAGCTTGACGGGGAGTTGCAGCAAGCGGTCATTGGGCG |
| C_46 | GTTAAAATTCGCATTAATGTGAGCGAGTAACACACGTTGG |
| C_47 | TGTAGATGGGTGCCGGAAACCAGGAACGCCAG |
| C_48 | GGTTTTCCATGGTCATAGCTGTTTGAGAGGCG |
| C_49 | GTTTGCGTCACGCTGGTTTGCCCCAAGGGAGCCCCCGATT |
| C_50 | GGATAGGTACCCGTCGGATTCTCCTAAACGTTAATATTTT |
| C_51 | AGTTGGGTCAAAGCGCCATTCGCCCCGTAATG |
| C_52 | CGCGCGGGCCTGTGTGAAATTGTTGGCGATTA |
| C_53 | CTAAATCGGAACCCTAAGCAGGCGAAAATCCTTCGGCCAA |
| C_54 | CGGCGGATTGAATTCAGGCTGCGCAACGGGGGATG |
| C_55 | TGCTGCAAATCCGCTCACAATTCCCAGCTGCA |
| C_56 | TTAATGAAGTTTGATGGTGGTTCCGAGGTGCCGTAAAGCA |
| C_57 | TGGCGAAATGTTGGGAAGGGCGAT |
| C_58 | TGTCGTGCACACAACATACGAGCCACGCCAGC |
| C_59 | CAAGTTTTTTGGGGTCGAAATCGGCAAAATCCGGGAAACC |
| C_60 | TCTTCGCTATTGGAAGCATAAAGTGTATGCCCGCT |
| C_61 | TTCCAGTCCTTATAAATCAAAAGAGAACCATCACCCAAAT |
| C_62 | GCGCTCACAAGCCTGGGGTGCCTA |
| C_63 | CGATGGCCCACTACGTATAGCCCGAGATAGGGATTGCGTT |
| C_64 | AACTCACATTATTGAGTGTTGTTCCAGAAACCGTCTATCAGGG |
| C_65 | ACGTGGACTCCAACGTCAAAGGGCGAATTTGGAACAAGAGTCC |
